# Role of the G-Protein Coupled Receptor 3-Salt Inducible Kinase 2 Pathway in Human β Cell Proliferation

**DOI:** 10.1101/2021.05.31.446443

**Authors:** Caterina Iorio, Jillian L. Rourke, Lisa Wells, Jun-Ichi Sakamaki, Emily Moon, Queenie Hu, Tatsuya Kin, Robert A. Screaton

## Abstract

Loss of pancreatic β cells is the hallmark of type 1 diabetes (T1D) ^1^, for which provision of insulin is the standard of care. While regenerative and stem cell therapies hold the promise of generating single-source or host-matched tissue to obviate immune-mediated complications^2–4^, these will still require surgical intervention and immunosuppression. Thus, methods that harness the innate capacity of β-cells to proliferate to increase β cell mass in vivo are considered vital for future T1D treatment^5, 6^. However, early in life β cells enter what appears to be a permanent state of quiescence ^7–10^, directed by an evolutionarily selected genetic program that establishes a β cell mass setpoint to guard against development of fatal endocrine tumours. Here we report the development of a high-throughput RNAi screening approach to identify upstream pathways that regulate adult human β cell quiescence and demonstrate in a screen of the GPCRome that silencing G-protein coupled receptor 3 (GPR3) leads to human pancreatic β cell proliferation. Loss of GPR3 leads to activation of Salt Inducible Kinase 2 (SIK2), which is necessary and sufficient to drive cell cycle entry, increase β cell mass, and enhance insulin secretion in mice. Taken together, targeting the GPR3-SIK2 pathway represents a novel avenue to stimulate the regeneration of β cells.

## Introduction

Progress in understanding human *β*cell proliferative behaviour has been hampered by the human *β* cell being a comparatively intractable system due to scarcity of tissue and a lack of genetic means to study them. Consequently, our knowledge of mechanisms of *β* cell proliferation comes largely from studies in rodents, where adult *β* cell proliferation is readily observed. As *β*cells are among the longest-lived cells in the mouse ^11^, mouse and human *β* cells share at least some degree of proliferative intransigence. However, key species differences in the cell cycle proteome ^8, 9, 12–14^ have necessitated research into human *β* cell biology. While phenotypic HTS of small molecule libraries is an efficient way to identify lead compounds that elicit human *β*cell proliferation to meet translational goals, a lack of understanding of the target and the mechanism of action of small molecule mitogens, as well as their off-target effects, remain a major impediment ^15^. We have argued that a safe regenerative approach will first require a genetic dissection of complex regulatory mechanisms governing the stable quiescence of human adult *β* cells ^16^; understanding the genetic program that establishes and maintains this quiescence is of both significant biological interest and therapeutic promise. In a small-scale RNAi screen targeting cell cycle components, we previously reported that ∼10% of human *β* cells enter the cell cycle following silencing of the cyclin-dependent kinase inhibitors (CDKIs) CDKN2C/p18 or CDKN1A/p21 ^16^.

To make our lentiviral silencing approach scalable for genome-wide screens while remaining compatible with screening in primary human tissue (Figure 1a), we developed a robotics-compatible HTS protocol, starting from shDNA plasmid isolation through to virus purification and target cell infection. This approach generated lentivirus achieving >90% infection of adult human *β* cells (Figure 1b), which maintain their identity following infection over the 10-day assay time course (Supplementary Figure 1). We observed effective reduction of several target proteins with HTS virus, including PDX1, PTEN, p21, p18, and BCL-XL (Figure 1c). When PTEN was silenced, we observed an increase in phosphorylation of the PTEN target AKT at Ser473 (Figure 1d), confirming knockdown and that expected functional consequences are preserved following infection with HTS virus. For over 40 years, cell cycle analysis has relied upon DNA tumour viruses such as adenovirus and SV40 to identify critical cell cycle components, including the tumour suppressors p53 and pRb ^17^. Thus, we primed *β* cells to proliferate using SV40 large T Antigen (TAg) and stained for incorporation of the S-phase marker EdU in C-peptide+ cells (Figure 1e). Following silencing of control targets p18 or p21 with HTS virus, we observed a 3-fold increase in EdU+ C-peptide+ cells over non-targeting (NT) shRNA (Figure 1f), an effect we confirmed by staining for Ki67 (not shown). Under these conditions, <5% of C-peptide+ cells stain for activated caspase-3, and EdU+ cells are not caspase-3-positive (Supplementary Figure 2).

**Figure 1.**
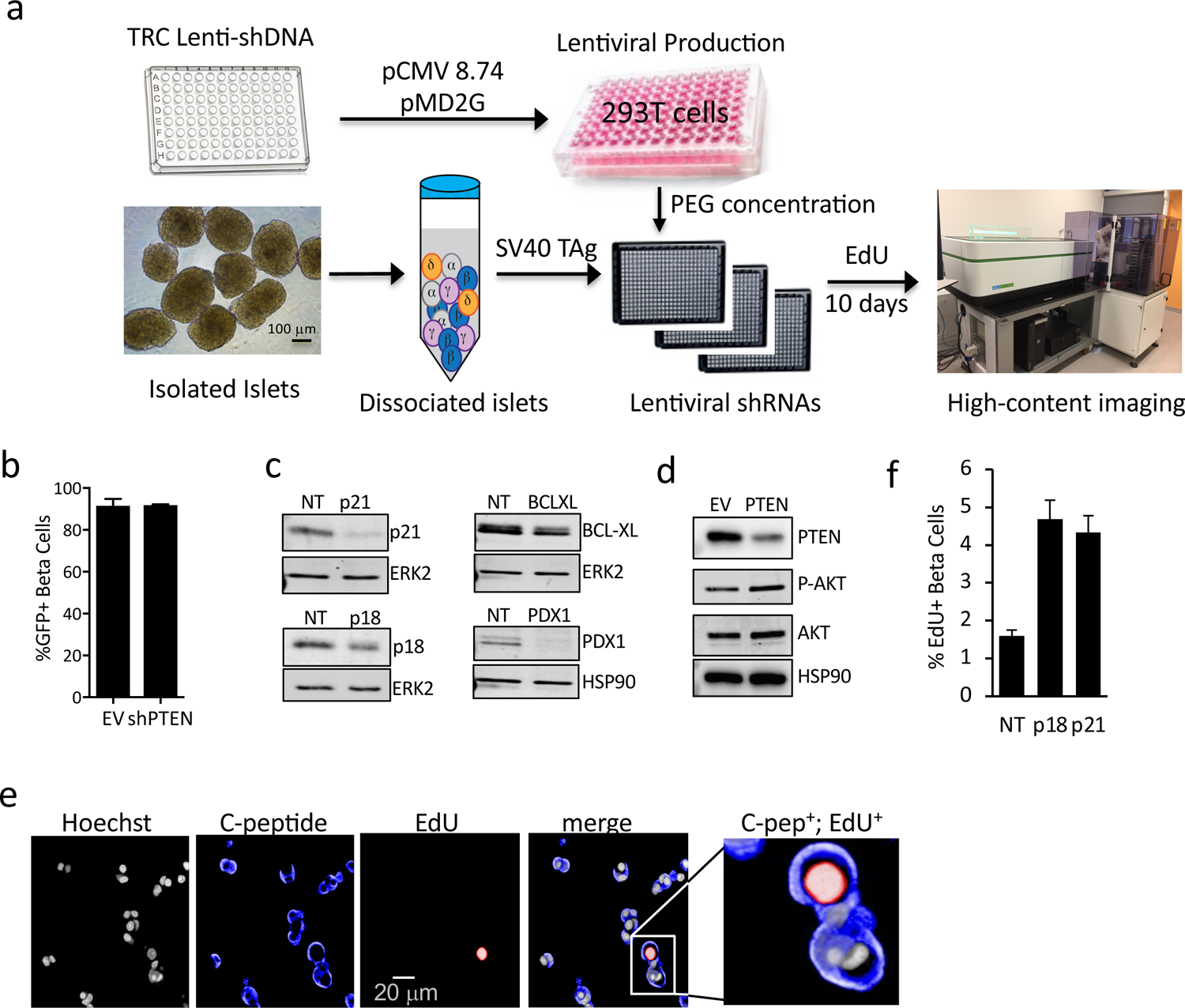
Automated HTS assay to screen in primary human cells. a. Schematic showing workflow of HTS screen, from shDNA plasmid packaging into lentivirus, arraying into 384-well plates and seeding with dissociated human islet cells, prior to 10d culture to permit gene silencing and EdU incorporation. b. Barplot showing % GFP positive, C-peptide positive cells after infection with HTS virus encoding shRNA targeting PTEN or empty vector control. c. Representative Western blots showing knockdown efficiency of positive controls following infection with lentiviral-shRNAs against p18, p21, BCL-XL, and PDX1. d. Effect of silencing PTEN on downstream effector phospho-AKT, with total AKT and HSP90 control is shown. e. Representative images of dissociated human islets from positive control well coinfected with TAg and silenced for p21, then stained for nuclei (Draq5, grey), C-peptide (blue), EdU (red), and merged. Scale bar = 20 um. Zoom merged image shown at right. f. Bar plot confirming effectiveness of lentivirus prepared with HTS protocol on silencing p18 and p21 and stimulating human β cell proliferation.

Human islets express numerous G protein-coupled receptors (GPCRs) ^18, 19^ which govern a myriad of key islet functions from hormone production and secretion to proliferation ^20–22^, and are important targets for diabetes treatment ^23–25^. The relatively low abundance of GPCR mRNA and protein permit efficient knockdown for RNAi screens and their druggability make them attractive targets for translation ^26, 27^. Therefore, we elected to silence 397 GPCRs and 40 of their cytoplasmic adaptors, the ‘GPCRome’, to identify GPCR signaling pathways that promote human *β* cell quiescence (see Table 1 for list of genes and shRNA sequences). We pooled 450,000 freshly isolated glucose-responsive islet equivalents from two non-diabetic donors (Supplementary Figure 3). Following dissociation, islet cells were seeded in 384-well plates containing 2342 pre-arrayed lentivirus encoding SV40 TAg together with individual library lenti-shRNAs, then cultured for 10 days in the presence of EdU. Z-scores for both the absolute number and percentage of C-peptide+ cells that also stained positive for EdU were determined (Figure 2a).

**Figure 2:**
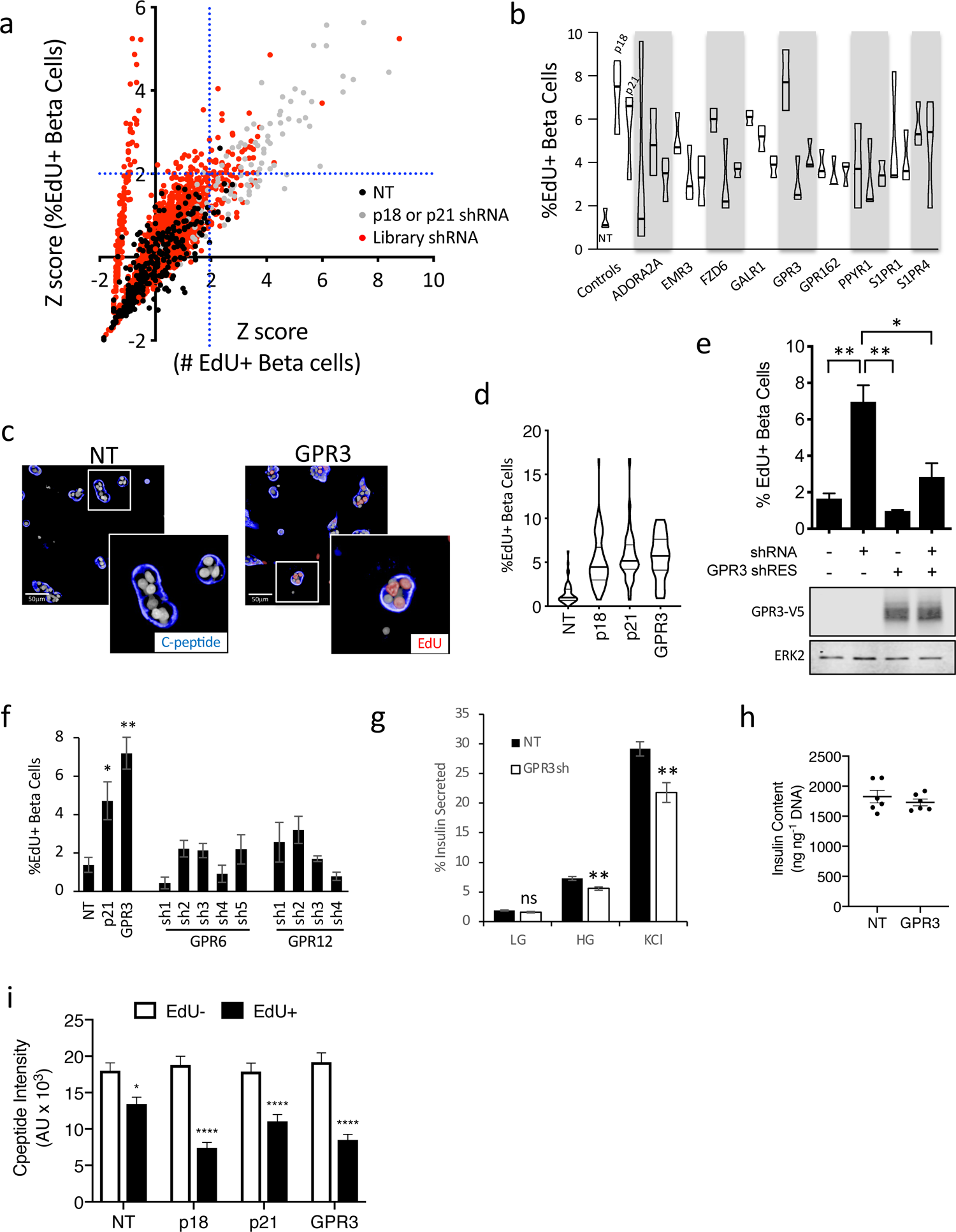
GPCRome screen identifies GPR3 as a regulator of human β cell proliferation. a. Z-score plots of screen results. Control cells infected with non-targeting virus (black circles, NT), positive control p18 or p21 shRNA virus (grey circles) or GPCRome library shRNAs (red circles) are shown. The x-axis represents Z-scores of double-positive (EdU, insulin) cells based on the absolute number; the y-axis represents Z-scores for the percentage of total β cells. Dotted blue lines mark the threshold of Z-scores = 2. b. Validation data for candidates identified in the screen. Percentage proliferation achieved for individual shRNAs against indicated targets were averaged over 3 screens using islets from 3 independent donors. c. Screen images showing representative NT control or GPR3 shRNA-infected β cells (C-peptide, blue) stained for EdU (red). d. Violin frequency distribution plots showing percentage of EdU+ insulin+ cells following GPR3 silencing in 18 donors over 10-day assay period compared to non-targeting control (NT) and p18 and p21 positive controls. Dark line = median, light line = quartiles. e. Barplot showing percentage of EdU+ β cells following silencing GPR3 in the presence of the shRNA-resistant GPR3 cDNA tagged with V5 (GPR3-RES-V5) or with NT shRNA control. Western blots showing expression of exogenous GPR3 is shown underneath. f. Barplot showing percentage of EdU+ beta cells following silencing of GPR3 and 5 independent shRNAs targeting GPR6 and 4 independent shRNAs targeting GPR12, the most closely related GPCRs to GPR3. g. GSIS in cells silenced for GPR3 compared to NT control. LG = low glucose, HG = high glucose, KCl = depolarizing stimulus. h. Insulin content in cells silenced for GPR3 compared to NT control. i. Barplot showing C-peptide intensity in EdU+ and EdU-populations in cultures silenced for indicated targets. Non-targeting (NT) control shown. All cultures were infected with TAg.

Nine candidates with at least two shRNAs with Z-scores ≥2 for both criteria were evaluated in confirmation screens using islets from three independent donors (Figure 2b). Of the candidates identified, G-protein-coupled receptor 3 (GPR3) has been implicated in cell cycle arrest and survival signaling in cerebellar development ^28, 29^, and in germ cell cycle arrest in *X. Laevis* ^30^ and mammalian oocytes ^31–33^, so we selected GPR3 for validation. Visual analysis of GPR3 silenced cultures revealed the presence of numerous EdU+ cells and even nuclear doublets (Figure 2c), suggestive of mitotic events. Silencing GPR3 mRNA (Supplementary Figure 4) gave significant increases in EdU incorporation in *β* cells in all subsequent donors tested (n = 18, Figure 2d). To rule out off-target effects of the GPR3 shRNA, we expressed an shRNA-resistant GPR3 cDNA to reconstitute GPR3 protein, which restored quiescence in the presence of GPR3 shRNA and reduced basal proliferation when expressed with control shRNA (Figure 2e). Silencing GPR3’s closest phylogenetic neighbours, GPR6 and GPR12 ^34^, did not induce proliferation above baseline, indicating a specific role for GPR3 in maintaining cell cycle arrest (Figure 2f). The silencing of GPR3 also increased *β*cell proliferation in the absence of TAg (Supplementary Figure 5). Taken together, we conclude that silencing GPR3 can reverse the stable quiescence of adult human *β*cells. Knockdown of GPR3 in dispersed and intact islets had a negligible effect on glucose-stimulated insulin secretion (GSIS) (Figure 2g) or insulin content (Figure 2h), showing *β* cells lacking GPR3 maintain functionality. In contrast, the intensity of C-peptide staining in EdU+ cells was 25-70% lower than in EdU-cells (Figure 2i), consistent with previous observations documenting downregulation of insulin expression in proliferating cells^16^.

We previously reported that silencing the CDKIs p18 and p21 in human *β* cells leads to S-phase entry ^16^. We reasoned that silencing GPR3 would reduce levels of one or more CDKIs, so we performed Western blots for the INK4 class (p15, p16, p18) and the more broadly acting CIP/KIP class (p21, p27, and p57) of CDKIs. Of these, only p21 and p27 of the CIP/KIP class of CDKIs were reduced following GPR3 silencing (Figure 3a and 3b); the CIP/KIPs CDKIs inhibit the assembly and catalytic activity of Cyclin D-dependent CDK complexes in the G0-G1 transition, Cyclin E-CDK2 complexes in G1, and Cyclin A-CDK2 complexes in the S to G2 transition ^35^. The CIP/KIP class are turned over by a SCF ubiquitin ligase complex that harbours the F-box protein SKP2 as its substrate-selectivity component ^36^, raising the possibility that GPR3 signaling may reduce SKP2 levels, and in turn, increase levels of CIP/KIPs. Unexpectedly, SKP2 levels were not increased following GPR3 silencing, indicating that accumulation of SKP2 does not mediate the loss of CIP/KIP proteins in this context (Figure 3a). Consistent with our previous work, silencing p21 induced *β*cell cycle entry (Figure 3c). To determine if loss of p27 or p57 alone is sufficient for the induction of proliferation, we silenced each with shRNAs and observed proliferation following p27 ^37^ (Figure 3d), but not p57 ^38^ (Figure 3e) silencing. To determine if GPR3 and the CDKIs p18^39^, p21 and p27 work in the same pathway, we performed co-silencing experiments to explore possible epistatic relationships between GPR3 and the CDKIs (Figure 3f). Whereas co-silencing p18 or p21 with GPR3 resulted in enhanced proliferation over the CDKIs alone, silencing GPR3 together with p27 did not. Taken together, our data suggest that p18, p21 and p27 are the key CDKIs that prevent proliferative activity in adult human *β*cells, and that GPR3 promotes cell cycle arrest by maintaining levels of p21 and p27.

**Figure 3:**
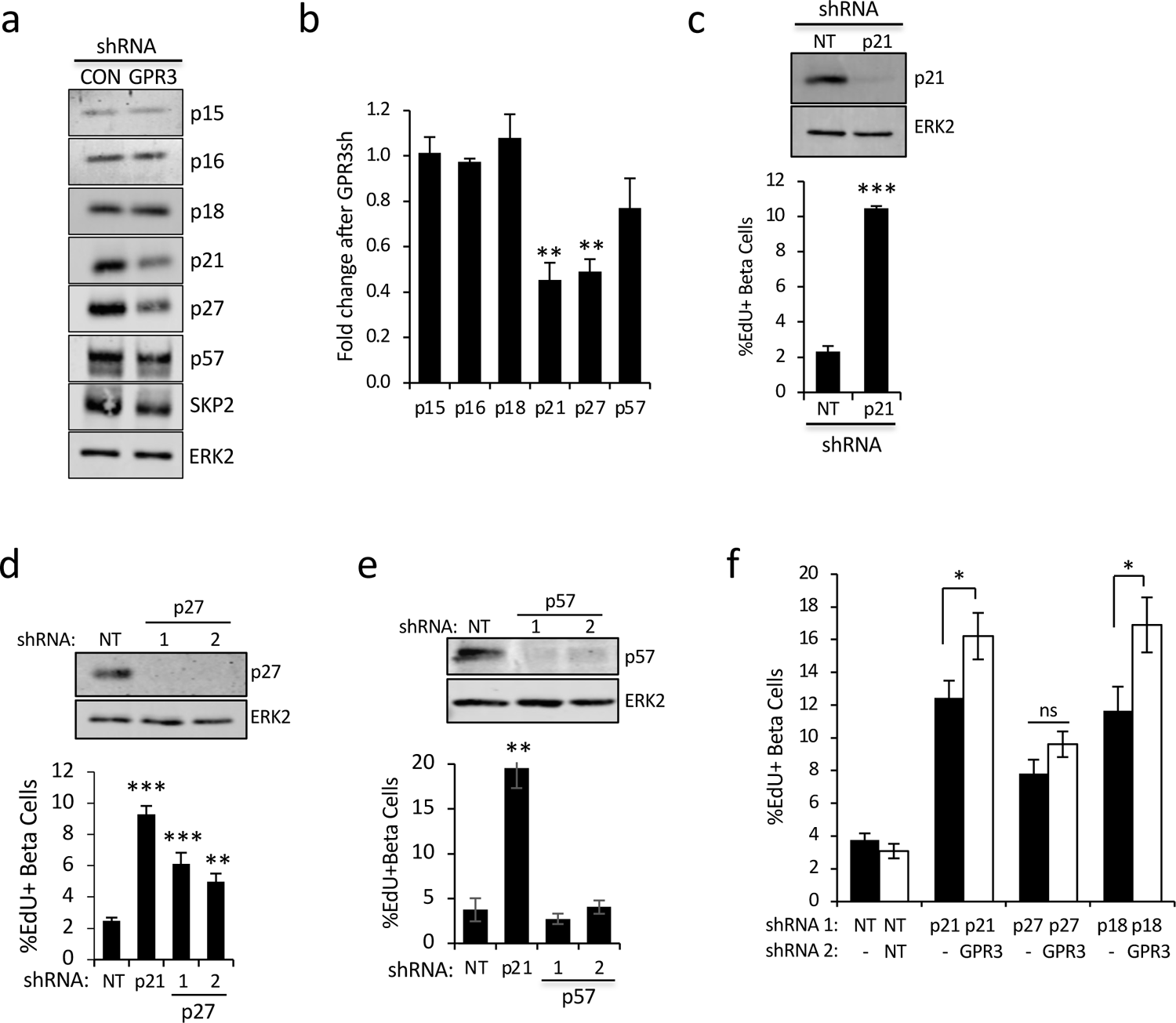
GPR3 stabilizes CIP/KIP cell cycle dependent kinase inhibitors. a. Representative Western blot analysis of INK and CIP/KIP family CKI levels in NT control and GPR3-silenced human islet cells. b. Barplot showing quantitation of CKI intensities. c-e. Western blots and barplots of human beta cell proliferation following silencing of p21 and p27, and p57. All samples co-express TAg. f. Barplot showing human beta cell proliferation following silencing of p18, p21, or p27 alone or in combination with GPR3. All samples co-express TAg. No virus added is indicated by a dash. NT = non-targeting shRNA control.

### The GPR3-SIK pathway maintains human β cell quiescence

GPR3 promotes constitutive cAMP synthesis through adenylyl cyclase via an unknown ligand ^31^ which prompted us to evaluate the role of PKA signaling in GPR3-mediated cell cycle arrest ^40, 41^. We monitored the phosphorylation status of the transcriptional coactivator CRTC2, which becomes dephosphorylated when PKA is active ^40^. Western blots of GPR3 silenced cells show that CRTC2 became more phosphorylated, consistent with inhibition of CRTC2 and reduced PKA activity (Figure 4a). As Ser275 on CRTC2 is a target of Salt Inducible Kinase 2 (SIK2) ^41, 42^, we treated islets with the pan-SIK inhibitor HG-9-91-01 (SIK-in), which led to CRTC2 dephosphorylation (Figure 4b) and restored quiescence in GPR3-silenced cells (Figure 4c). Taken together, we conclude that GPR3 promotes quiescence in the adult *β* cell by inhibiting SIKs.

**Figure 4.**
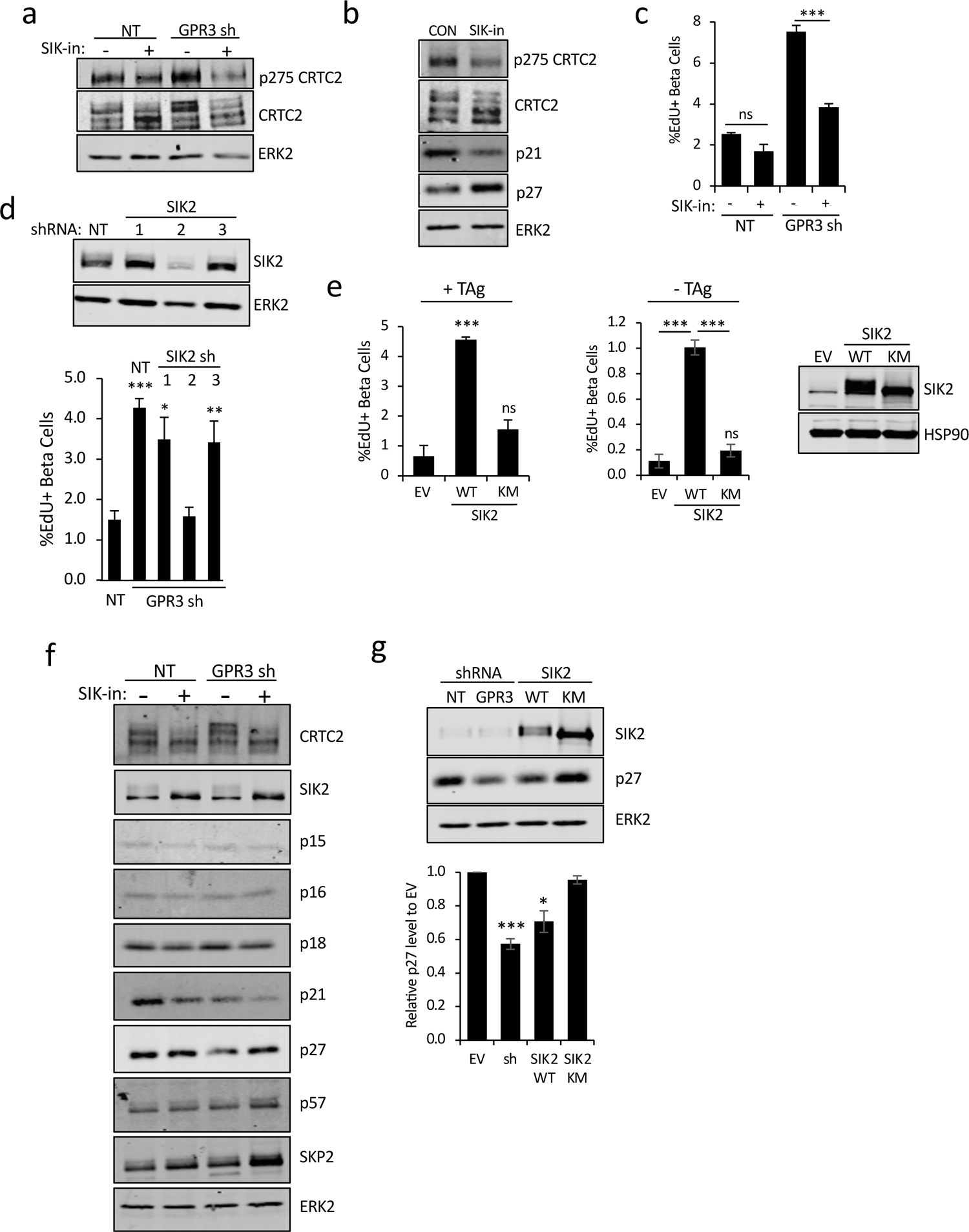
GPR3 promotes quiescence by inhibiting SIK2. a. Western blot showing increase in phosphorylation status of SIK2 target protein CRTC2. Effect of pan-SIK inhibitor (SIK-in) is shown. b. Western blot showing effect of SIK-in on levels of p21 and p27 in human islet cells. c. Barplot showing effect of SIK-in on proliferation in GPR3-silenced human beta cells. d. Effect of co-silencing SIK2 in cells silenced for GPR3 on beta cell proliferation. Western blots showing silencing shown. e. Barplots showing effect of overexpression of SIK2 WT and kinase dead (KM) mutants of SIK2 on beta cell proliferation in the presence of TAg (left) and absence of TAg (middle). EV vs. KM = ns. (right) Western blot showing levels of SIK2 proteins shown. f. Western blots showing levels of CIP/KIP proteins in GPR3 silenced beta cells. Effect of pan-SIK inhibitor HG-9-91-01 (SIK-in) shown. g. Western blots showing p27 levels in cells following GPR3 silencing or overexpression of SIK2 WT or kinase dead mutant (KM). Barplot showing quantification (n=3) shown at bottom.

As SIK-in inhibits SIK1, SIK2 and SIK3, we next used RNAi to identify which SIKs are required for GPR3 to maintain *β* cell quiescence. When GPR3 was silenced together with each SIK, proliferation rates reversed back to control levels only when SIK2 was co-silenced with GPR3 (Figure 4d), silencing SIK1 and SIK3 had no effect (Supplementary Figure 6). Consistent with a requirement for SIK2 for *β* cell proliferation, overexpression of a wild type SIK2, but not a kinase-dead SIK2 mutant (KM), was able to increase proliferation rates to those observed in GPR3-silenced cells both in the presence and absence of TAg (Figure 4e). To determine which CDKIs are regulated by SIKs, we treated control and GPR3-silenced cultures with SIK-in and performed Western blots. Whereas levels of p21 and p27 are reduced by GPR3 silencing, only levels of p27 protein were restored following SIK inhibition (Figure 4f). Consistent with this, overexpression of SIK2 in human islets, but not SIK2 KM, reduced levels of p27 as effectively as silencing GPR3 (Figure 4g).

Deregulated SIK2 has been reported to support proliferative behaviour in other contexts ^43^, thus we postulated that enhanced levels of SIK2 may enhance proliferation of *β* cells *in vivo*. To address this, we generated SIK2 (MIP-SIK2-V5) transgenic (Tg) founder mice that expressed ∼2-4-fold higher SIK2 in isolated islets than control mice (Figure 5a; Supplementary Figure 7), levels similar to those we observed in non-diabetic ob/ob or Western diet-fed mice ^41^. Ki67 staining (Figure 5b) revealed that elevated SIK2 expression was sufficient to increase average proliferation rates from 0.3% to over 2% (Figure 5c) and increase islet area 2-fold (Figure 5d). At 16-20 weeks, MIP-SIK2 Tg mice display normal fasting blood glucose and normal glucose tolerance (Supplementary Figure 8) but reduced blood glucose levels after 1 hr refeeding (Figure 5e). Interestingly, these SIK2 Tg mice secrete significantly higher basal insulin secretion as well as after arginine challenge (Figure 5g), characteristic of a fetal beta cell functional profile ^44^.

**Figure 5:**
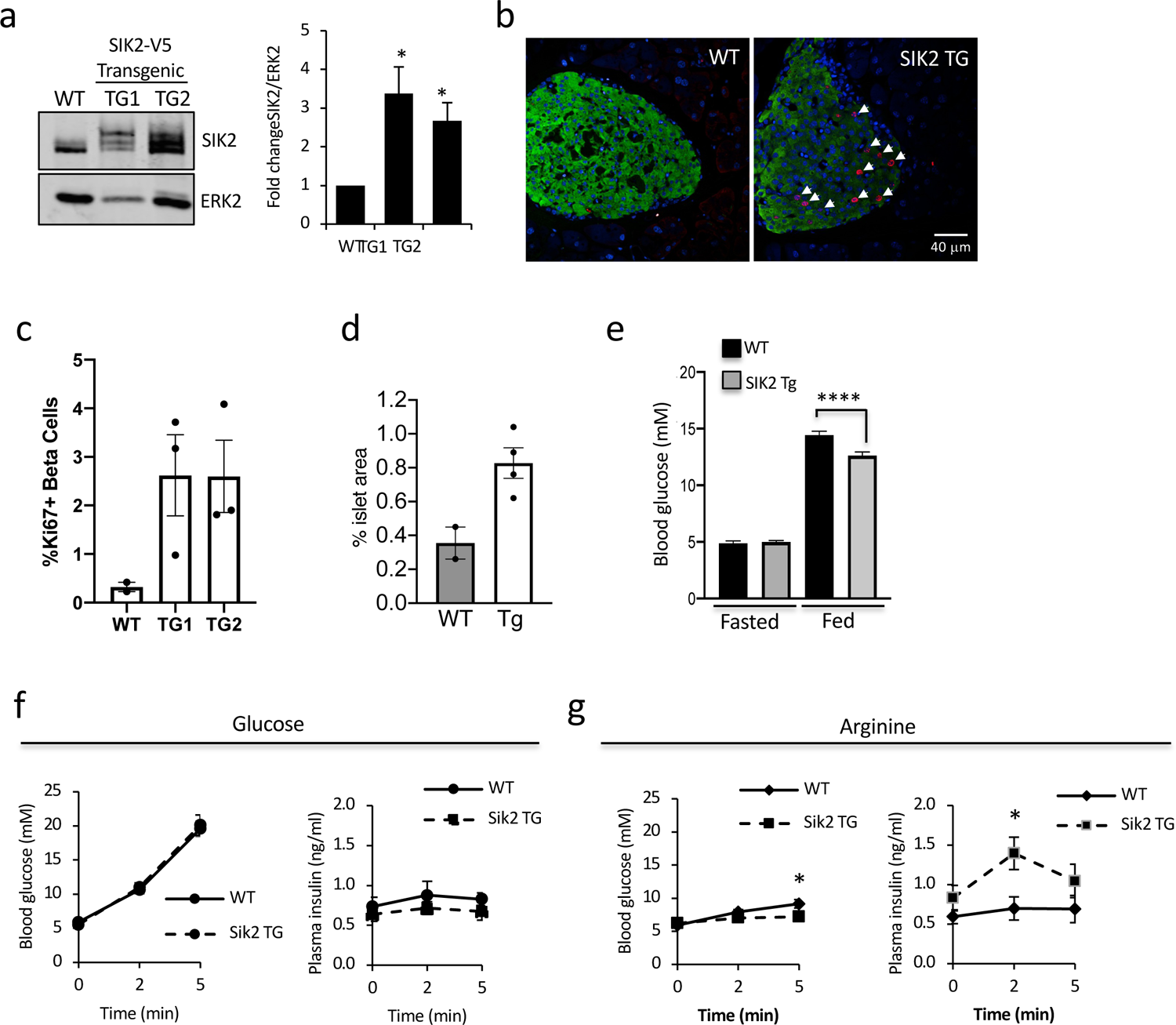
SIK2 promotes mouse beta cell proliferation. a. (left) Western blot showing levels of endogenous and SIK2-V5 protein in control and MIP-SIK2 transgenic (SIK2-V5-TG) animals with ERK2 loading control. (right) Barplot showing fold change in SIK2 protein in SIK2 TG1 and TG2 relative to WT (10-14 weeks, n=3). b. Ki67 (red) immunostaining of pancreatic sections from WT (left) and SIK2 Tg mice (12-16 weeks). Arrowheads show proliferating beta cells. Hoechst staining of nuclei shown (blue). c. Barplot showing quantification of Ki67+ insulin+ cells from WT and SIK2 transgenic founders 1 and 2. 1000-1500 cells were counted for each. d. Barplot showing % islet area in WT and SIK2 Tg mice. e. Barplot showing fasting and fed blood glucose concentrations for WT and founder line 2 of MIP-SIK2 (SIK2 Tg) mice at 16 weeks of age (n = 10 WT, 15 Tg). f,g. Line charts showing blood glucose (left) and plasma insulin (right) in WT (solid line) and SIK2 Tg (dotted line) mice (8-16-weeks) following IP injection of (f) glucose or (g) arginine.

Consistent with these data, SIK2 protein in islets isolated from non-diabetic human subjects ranging from lean to obese BMI (Figure 6a) increased linearly (Figure 6b, r^2^ = 0.52), supporting the notion that increased SIK2 in the human islet permits a compensatory secretory response. Levels of p27 were reduced with increasing BMI, indicating that increasing levels of SIK2 correlates with loss of p27 in human subjects (Figure 6c-d). Levels of the *β* cell marker PDX1 were unchanged (Supplementary Figure 9). We previously described a complex consisting of SIK2, the CDK5 activator CDK5R1/p35, and the E3 ligase PJA2 that is essential for functional compensation in the *β* cell, in which SIK2 protein accumulates in islets of prediabetic mice^41^. In keeping with our animal data, silencing SIK2 in human islets resulted in increased levels of the SIK2 target p35 (Figure 6e) and reduced glucose stimulated insulin secretion (GSIS) by 35% without affecting insulin content (Figure 6f). Similarly, treatment with SIK-in to inhibit all SIKs reduced GSIS 40% without affecting insulin content (Figure 6g), consistent with a specific role for SIK2 in this regard.

**Figure 6.**
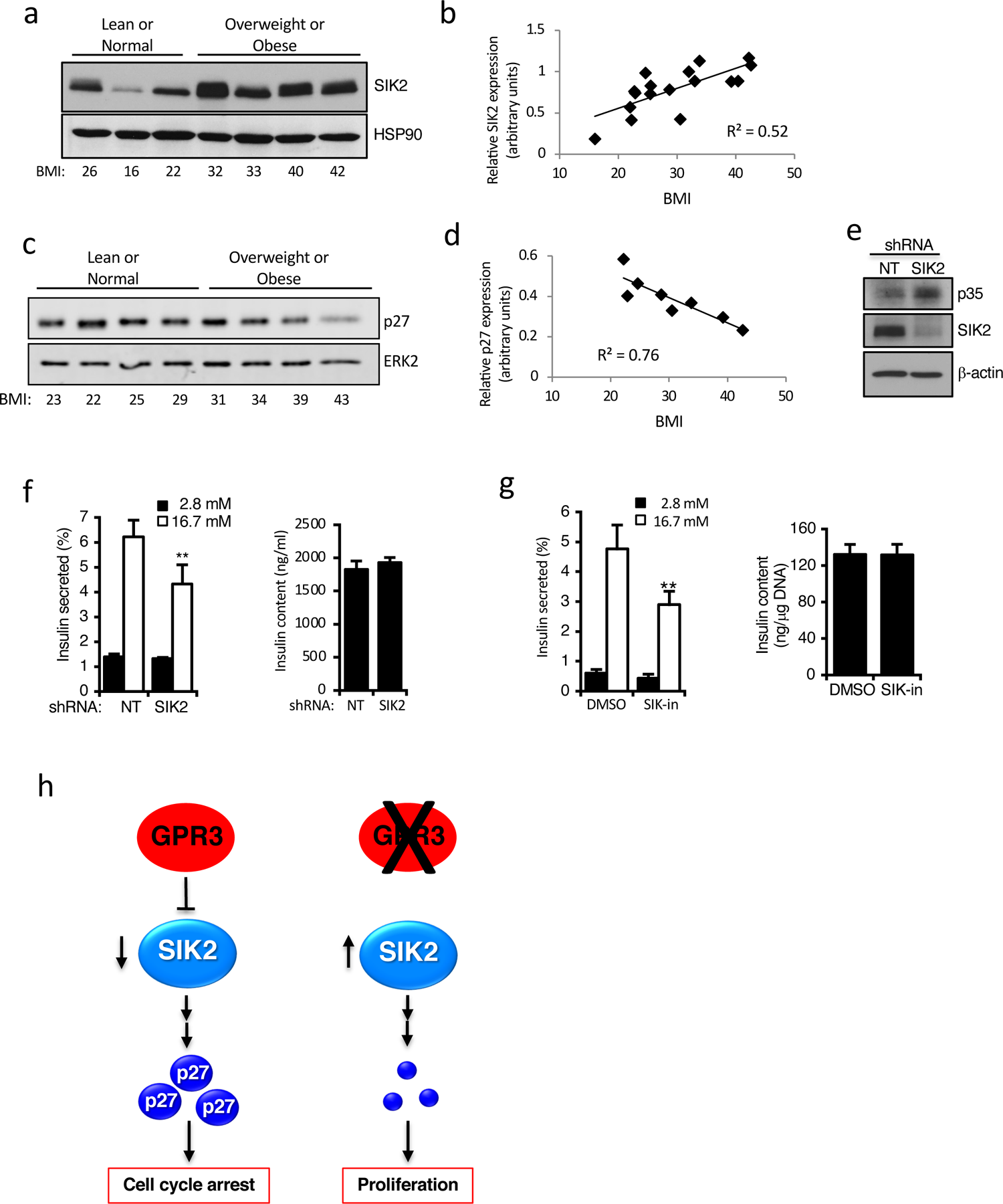
SIK2 promotes human beta cell function. a. Western blots showing levels of SIK2 protein and HSP90 loading control in isolated human islets from non-diabetic human subjects of increasing BMI. b. Line plot showing correlation between SIK2 levels in isolated human islets and subject’s BMI; r^2^ = 0.52. c. Western blots showing levels of p27 protein and ERK2 loading control in isolated human islets from non-diabetic human subjects of increasing BMI. d. Line plot showing correlation between SIK2 levels in isolated human islets and subject’s BMI; r^2^ = 0.76. e. Western blot showing levels of SIK2 and its substrate p35 following silencing of SIK2 compared to NT control. f. Barplots showing insulin secretion in response to 16.7 mM glucose (left) and insulin content (right) following silencing of SIK2 in isolated human islets compared to NT control. g. Barplot showing effect of SIK-in on GSIS (left) and insulin content (right) in isolated human islets compared to DMSO control. h. Model showing GPR3 promoting p27 accumulation to maintain cell cycle arrest. When GPR3 is silenced, SIK2 becomes active and p27 is lost, facilitating cell cycle entry.

## Discussion

Thus, strategies designed to restore lost or damaged tissue must be subject to reversal and will require a complete understanding of the proliferative machinery and their regulation in the human *β* cell. Our work identifies GPR3 as a key cell surface receptor that suppresses SIK2 activity to stabilize CIP/KIP CDKIs and maintain quiescence. We found that silencing GPR3 in the presence of TAg triggered ∼5-6% of primary human β cells to proliferate, which could be rescued by restoration of GPR3 with RNAi-resistant cDNA, establishing GPR3 as a *bona fide* target.

Consistent with this, mild overexpression of GPR3 reduced proliferation in control cells, indicating that proliferative potential in human *β* cells is finely tuned to the levels of GPR3. That we do not achieve 100% proliferation with silencing p18, p21, p27, or GPR3 in the presence of TAg is consistent with our proposal that a complex genetic program consisting of multiple redundant anti-proliferative pathways is activated in the adult *β* cell to maintain a stable quiescence ^16^. Of note, GPR3 silencing lowers the more promiscuous CIP/KIP CDKIs, demonstrating its key role as a regulator of *β*cell quiescence and underscoring the likelihood that numerous roadblocks to *β* cell cycle entry must be overcome prior to entering S-phase. Owing to the ontological commonality between neurons and *β* cells, it is interesting that GPR3 also establishes quiescence of cerebellar granule neurons during postnatal development in mice ^28^, raising the possibility of a general role for GPR3 in regulating cell cycle entry decisions in excitable cells. Indeed, silencing GPR3 also resulted in cell cycle entry in *α* and *δ* cells, suggesting it can serve a broader role in suppressing cell cycle entry of the neuroendocrine lineage (Supplementary Figure 10). The GPR3 whole animal knockout results in age-dependent obesity and mild glucose intolerance due to impaired thermogenesis, but serum insulin and C-peptide levels were unchanged^45^. Extra-islet compensatory effects (brain, fat) in the GPR3 knockout will require clarification of GPR3’s role *in vivo* by tissue-specific deletion of GPR3. It is noteworthy that EdU-positive cells in all conditions show a reduction in C-peptide staining intensity, suggestive of loss of *β* cell functional identity in proliferating *β* cells^16^. This is in keeping with the idea that the fully mature *β* cell state and cell cycle entry are mutually exclusive, perhaps due to an energetic insufficiency preventing these programs from coexisting.

Consistent with a role for GPR3 as a constitutively active Gs-coupled GPCR, silencing GPR3 leads to hyperphosphorylation of CRTC2, a PKA-SIK axis substrate, suggestive of activation of CREB-dependent transcription ^40^. However, we see no evidence of steady-state activation of CREB targets in cells lacking GPR3 (data not shown). Treatment of human islets cells lacking GPR3 with pan-SIK inhibitor prevents hyperphosphorylation of CRTC2, implicating SIKs as critical effectors downstream of GPR3. While our data support a GPR3-SIK2-p27 pathway as central to regulating *β* cell quiescence, silencing GPR3 also reduced levels p21 which were not restored by inhibiting SIKs, pointing to the existence of additional effectors downstream of GPR3 that mediate p21 status. However, as silencing SIK2 in cells lacking GPR3 restored quiescence and overexpression of SIK2 in the absence of TAg in human *β* cells promoted proliferation *in vitro* and *in vivo*, activation of SIK2 alone is necessary and sufficient for cell cycle entry. *β* cell knockout of SIK2 results in impaired glucose-stimulated insulin secretion via reduced phosphorylation of p35 and inactivation of voltage-gated calcium channels, without a change in *β* cell mass^41^, suggesting compensation from SIK1 and SIK3 in this setting. Taken together, we propose that fluctuating SIK2 levels, dictated by ambient glucose concentrations, are a key element of the homeostatic process that fine tunes the insulin secretory response to meet demand. Should hyperglycemic conditions persist, sustained increases in SIK2 levels lowers the threshold for cell cycle entry via a reduction in p27 (Figure 6h). During this mass expansion, SIK2 Tg mice increase their responsiveness to amino acids, consistent with the functional profile of fetal beta cells ^44^ and stem cell derived *β*-like cells ^46^. Our data indicate that a successful regenerative approach to increasing functional *β* cell mass will involve the pharmacologic management of both cell cycle entry and of maturation.

Chemical biology approaches have identified candidate small molecule activators of *β* cell proliferation, including harmine, a pan-kinase inhibitor that induces 1% of human *β* cells to enter the cell cycle ^15^, by a mechanism that may involve as many as 14 kinases in the CMGC branch of the kinome ^47^. Silencing harmine in cells increase proliferation by 1.8% but had no effect when added to cultures silenced for GPR3 in the context of TAg expression (20% vs 21% when harmine added, Supplementary Figure 11). Harmine also reversibly inhibits monoamine oxidase A, and has been evaluated as a mood-altering therapy in human subjects ^48^. As several brain regions in GPR3 KO mice showed reduced levels of monoamine neurotransmitters such as serotonin and dopamine ^49^, the connection between GPR3 and the regulation of monoamine oxidases and their substrates warrants further investigation. We contend that identification and validation of gene targets that establish and maintain *β* cell quiescence will require a genetic approach, which will in turn provide a comprehensive functional framework to design strategies that allow precise control over their proliferative and functional behaviour.

Taken together, our work demonstrates the promise of functional genetic screens for dissecting therapeutically relevant state changes in primary human cells and demonstrate that GPR3 suppression of SIK2 activity is a key element of the network dedicated to repressing adult human *β* cell replication. As such, GPR3 and SIK2 represent novel targets for promoting proliferation of human *β* cells to increase *β* cell mass for the treatment of insulin insufficiency. We anticipate that validation of additional candidates from the screen will provide additional cues about pathways that govern adult human *β* cell quiescence, and broader application of RNAi screening in primary human cells to inform regenerative approaches designed to elicit expansion of other mature, non-dividing cell populations for therapeutic advantage.

## Methods

### Animals

All procedures involving mice were approved with the Animal Care Committee of Sunnybrook Research Institute. SIK2 *β* cell transgenic mice were prepared by injection of a linearized SIK2-V5 tagged cDNA under the control of the mouse insulin promoter (MIP-SIK2-V5) into pseudopregnant C57Bl6 blastocysts. Two founder lines were positive for SIK2-V5 protein in isolated islets by Western blotting, and these were expanded for analysis (Founder 1: Gene Targeting Facility, University of Connecticut, CT, USA; Founder 2: The Centre for Phenogenomics, Toronto, CA). Non-transgenic littermates were used as controls. For measurement of fasted and refed blood glucose levels, blood was analyzed using a OneTouch Ultra glucometer following a 16h fast with water, then again after 1h of refeeding with chow diet. Glucose tolerance and plasma insulin tests were performed as described ^41^.

### Antibodies/Reagents

Western Blot: Erk2 (Santa Cruz 1:1000), p21 (Cell Signaling 1:1000), p27 (Cell Signaling 1:1000), PTEN (Cell Signaling 1:1000), pAKT (Cell Signaling, 1:1,000), AKT (Cell Signaling 1:1,000), p18 (Abcam 1:2000), PDX1 (Cell Signaling 1:1000), BclXL (Cell Signaling 1:1000), HSP90 (Santa Cruz 1:1000), V5 (Cell Signaling 1:1000), SKP2 (Cell Signaling 1:1000), p57 (Cell Signaling 1:1000), p16 INK4a (Cell Signaling 1:1000), p15 INK4b (Abcam 1:1000) and SIK2 (Cell Signaling 1:1000). Immunofluorescence: insulin (Dako 1:1,000), PDX1 (Abcam AB47383 1:2000), cleaved caspase 3 (Cell Signaling 1:100). The C-peptide antibody was from the Developmental Studies Hybridoma Bank, created by the NICHD of the NIH and maintained at The University of Iowa, Department of Biology, Iowa City, IA 52242. Antibodies for CRTC2 and pSer275 CRTC2 were described previously^42^. Click-it EdU Alexa Fluor 647 imaging kit (Life Technology) was used according to the manufacturer’s recommendations. Harmine was purchased from Cayman Chemicals and GNF4877 was a gift from J. Annes. Plasmids are described in Table 4.

### Cell and islet culture

HEK293T-17 cell culture has been described^41^. We used four sources of human pancreatic islets for assay development: the NIDDK-funded Integrated Islet Distribution Program (IIDP islets); University of Alberta, Edmonton, Canada, Clinical Islet Lab of J. Shapiro (Shapiro islets), Alberta Diabetes Institute Research Islet Lab, Canada, lab of P. MacDonald (ADI IsletCore islets), and Toronto University Health Network group, M. Cattral. Table 2 provides complete donor information and the experiments for which individual subjects were used. Male and female deceased donors were used, ranging in age from 17-78 years old and BMI from 18-44.4 kg/m^2^, none of which had a prior diagnosis of diabetes. When islet purity was under 85%, islets were picked manually and cultured for up to 14 days at 37°C in a 5% CO_2_ atmosphere in non-tissue culture treated petri dishes in PIM(S) media supplemented with 5% human AB serum, glutamine/glutathione mixture and penicillin/streptomycin (all reagents from Prodo Laboratories Inc.). Medium was changed every 2-3 days.

### GSIS

Intact human islets (50 islets per replicate) or re-aggregated (5000 cells/ well in V-bottom 96-well plate) human islets were equilibrated in Krebs Ringer Buffer (KRB) containing 2.8 mM glucose for 30 min, then incubated in 2.8 mM glucose in KRB for 1 hr prior to stimulation with 16.7 mM glucose in KRB for 1 hr, followed by 45mM KCl for 1hour. KRB supernatants from 2.8 mM, 16.7 mM glucose and 45mM KCl treatments were collected and insulin amount determined using a human insulin HTRF assay (Cisbio). Cells infected with lentivirus were incubated for 6 days prior to GSIS assay. All conditions were done in triplicate or quadruplicate.

### Islet Dissociation and Seeding

Islets were washed in PBS and dissociated with Accutase (1 ml/1000 IEQ) for 5-7 min at 37^°^C and triturated every 60 sec. Dissociated islet cells were seeded at a density of 15,000 cells/well in a 384-well plate for fluorescence or 60,000 cells/well in a 96-well plate to generate protein extracts. Islets were always seeded on a PDL-coated plate to facilitate attachment of dissociated cells^16^.

### Lentiviral shRNA

plasmids were from the MISSION Human shRNA library (Millipore SIGMA). The pLKO.1eGFP control vectors (EV, NT, p18, p21) and pLenti6.3-SV40T Antigen-V5 (TAg), as well as all lentiviral purification by ultracentrifugation have been described ^16^.

### HTS plasmid purification

GPCR genes were identified using data from the IUPHAR GPCR list and UNIPROT data for known 7 transmembrane-spanning receptors (accessed November 2016). Known pseudogenes were excluded (Table 1). The resulting list of 437 genes consisted of 397 GPCRs and 40 GPCR-related and associated proteins. The shRNAs for 7 GPCRs and 2 GPCR-related proteins were unavailable in the Mission TRC library resulting in 98% coverage of the selected GPCRome and a total of 2342 shRNAs. 1-26 shRNAs (average = 5) were available per gene. Frozen glycerol stocks from the GPCR shRNA gene set were scraped, inoculated into 4 ml of TB medium containing 50 μg/ml ampicillin in a 24-well deep-well culture plate, and grown for 18 hr at 37°C with 250 rpm shaking. Bacteria were pelleted by centrifugation at 1000 x g for 10 min. Plasmids were isolated using 96 well NucleoSpin transfection-grade DNA kits (Machery-Nagel) measuring absorbance at 260 nm and 280 nm and verified by agarose gel electrophoresis.

DNA was diluted to 10 ng/ml in 5 mM Tris/HCl (pH 8.5) using a Mantis Liquid Handler (Formulatrix), arrayed and stored in a robotic −20C storage system (Hamilton Verso).

### HTS Lentiviral packaging

Lentiviral packaging in HEK293T-17 cells has been described ^16^. Viral packaging (pCMV8.74, 45 ng/well), envelope (pMD2G, 6 ng/well) and shRNA-containing transfer (pLKO.1-shRNA, 50 ng/well) plasmids were combined in 20 μl/ well OptiMEM (Life Technologies) containing 0.2 μg/well linear polyethyleneimine 25,000 (PEI) transfection reagent in a 96-well tissue culture plate. Following incubation for 15 min at room temperature, 10^5^ HEK293T-17 cells in 100 μl DMEM containing 10% FBS and 5 IU/ml penicillin/streptomycin were added to the DNA mixture, mixed by pipetting, and incubated at 37°C 5% CO_2_ for 72 hr. HEK293T-17 cells were maintained below 90% confluence for no more than 20 passages.

### Lentiviral purification

Cell supernatants were vacuum filtered (0.2 μm, 96-well - Agilent Technologies) and viral particles were mixed with polyethylene glycol 8000 (SIGMA) to a final concentration of 5-6% and incubated at 4°C for 5 hr. Lentivirus was pelleted by centrifugation at 2500 x g for 1 hr and resuspended in 30 μl PIM(S) medium. Purified lentivirus was arrayed in black 384-well poly-D-lysine-coated imaging plates (Greiner) and stored at −80°C until use ^16^.

### shRNA-resistant cDNA rescue

The open reading frame of GPR3 cDNA (Genscript) and SIK2 ^40^ was cloned into a modified version of pLenti6/V5-DEST vector (Invitrogen) in which we deleted 300 nucleotides of the CMV promoter distal to the transcription start site were removed to attenuate its potency as per ^50^. The resulting expression plasmid, pLenti6-delta4/V5-DEST provides more physiological expression levels appropriate for genetic rescue experiments. shRNA-resistant constructs were generated by QuickChange Lightning Site-Directed Mutagenesis Kit (Agilent). PCR oligo sequences and mutagenesis primer sequences are listed in Table 3.

### Islet cell infection

Dissociated islet cells were infected at the time of seeding by adding the cell suspension to multi-well plates (96 or 384-well) in which purified lentivirus had been arrayed. For high-potency concentrated lentivirus prepared by ultracentrifugation, ^16^ islet infection in 384 well plates was performed using 0.5% of the yield from a 10 cm dish for all viruses. For 96 well plates, 5% of the yield was used. For GPCR screen shRNAs, islets in 384-well plates were infected with 100% of yield from a 96 well dish of PEG-purified lentivirus. For epistasis experiments, one shRNA virus (p18sh, p21sh or p27sh) was added together with lenti-TAg at the time of seeding; 24 h later cells were rinsed twice with PIMS medium and the second virus (GPR3sh) was added. Harmine (10 uM) and GNF4877 (2 uM) were added to wells 3d and 7d after seeding prior to imaging on d10.

### Immunofluorescence

EdU was added to the medium at 10 μM on 3 and 7 days after seeding. Following 10 days of lentiviral infection, dissociated islet cells were fixed by adding 3.7% paraformaldehyde containing Hoechst (5 μg/ml) for 15 min at 37°C and quenched with an equal volume of 0.75% glycine in PBS for 5 min. Cells were permeabilized with 0.1% Triton-X-100 in PBS for 10 min. Click-IT EdU detection was performed according to the manufacturer’s protocol, prior to blocking in 3% BSA in PBS overnight. Antibodies in 3% BSA were incubated overnight at 4^°^C (primary) or for 1 hr at RT (secondary). Each step was followed by two gentle washes in PBS using a BioTek 405select multiwell plate washer.

### Imaging and Data Analysis

Fluorescent images were captured using a Perkin-Elmer Opera Phenix automated confocal multiwell plate microscope fitted with a 40X high NA water lens. 49 fields per well were acquired in each fluorescence emission channel: blue 405 (Hoechst), green 488 (GFP), red 594 (C-peptide/Insulin/PDX1) and far-red 647 (EdU). Columbus high-content imaging analysis software was used to generate an algorithm to identify the intensity of each stain in regions of interests in an object (cells) and to determine the % of nuclear EdU+ (proliferating) and C-peptide+ (insulin positive) *β* cells that were GFP+ (infected with lenti-shRNA virus).

### RT-PCR

Islets were dissociated as previously described and 400,000 cells were infected with NT or GPR3 shRNA lentivirus in a poly-D-lysine-coated 24 well plate. 6 days post-infection, cells were collected and lysed with TRIZOL (manufacturer) and total RNA was isolated according to the manufacturer’s protocol. Any contaminating DNA was removed from the sample using the DNA-free DNase Treatment and Removal Kit (Thermo Fisher Scientific) according to the manufacturer’s protocol. Total RNA was transcribed into cDNA using random primers using the Superscript III First-Strand Synthesis kit (Thermo Fisher Scientific) and amplified with PDX1 and GPR3-directed oligos described in Table 3 and Supp. Fig 4.

### SIK inhibitor treatment

Dissociated islets were plated on either 384-well plates (for imaging) or 96-well plates (for Western blots) and infected with the indicated lentivirus. 24 h after infection, cells were treated at a final concentration of 2 μM HG-9-91-01 (APExBIO) for duration of the experiment. Fresh drug was added daily.

### Western blotting

Six days after lentiviral infection, dissociated human islet cells were washed twice with ice-cold PBS prior to lysis in 2x Laemmli buffer. Sample proteins were separated by SDS-PAGE electrophoresis and transferred to low-fluorescence PVDF membrane (Millipore). Membranes were blocked for 1 hr in 3% milk in TBST prior to incubation with the indicated primary antibodies in 3% BSA overnight at 4°C. IRDye secondary antibodies (Licor) were diluted 1:30000 (IR680) or 1:18000 (IR800) in 3% milk in PBS containing 0.01% SDS and 0.04% TritonX-100 and incubated with membranes for 1 hr at room temperature. A Licor Odyssey Clx was used for signal detection. All Western blots are representative of at least n=3 experiments.

### β Cell Proliferation *in vivo*

Pancreases from 12-16 week old WT (n=2), SIK2 Tg founder 1 (n=3) and founder 2 (n=3) mice were isolated and fixed O/N in 4 % paraformaldehyde/PBS, then rinsed with 70% ethanol prior to paraffin embedding at the Histology Core at Sunnybrook Research Institute (SRI). Blocks were then sectioned and stained for Hoechst (blue), Insulin (Green) and Ki67 (Red) at the Histology Core at the University Health Network. The percentage of Ki67+ insulin+ cells in islets was determined from 3 stained sections per mouse approximately 150 μm apart. The stained sections were imaged using the Perkin-Elmer Opera Phenix with 40X high NA water lens.

### β cell area

Pancreases were sent to Histology Core at SRI for H&E stain and paraffin embedding. Pancreatic sections were taken 150 μm apart, and deparaffinized prior to antigen retrieval in 0.1M sodium citrate pH6 and staining with guinea pig anti-insulin antibody (Dako A0564, 1:250) overnight at 4 degrees in the dark and visualized with Anti-Guinea Pig biotin conjugated antibody (Vectastain BA7000, 1:300). Slides were imaged at the Advanced Optical Microscopy Facility, Toronto, and images analyzed using Aperio ImageScope.

### Statistical Analyses

were performed using GraphPad Prism® 8 or Excel Software. Statistical significance was determined using Student’s t-test when comparing two means; one-way ANOVA and Tukey post tests were applied for comparison between different groups. p-values of <0.05 were taken to indicate statistical significance (ns or no asterisk = not significant, p ≥ 0.05, * p ≤ 0.05, ** p ≤ 0.01, *** p ≤ 0.001, **** p ≤ 0.0001). Unless otherwise indicated, error bars represent +/- standard error of the mean across 3 independent donors/experiments.

## Acknowledgements

We acknowledge M. Cattral and the Cell Isolation Processing Facility (University Health Network, Toronto), the Integrated Islet Distribution Program (IIDP), and P. MacDonald and J. Manning-Fox of the ADI IsletCore for provision of human pancreatic islets. Islet isolation at the ADI IsletCore was subsidized in part by funding from the Alberta Diabetes Foundation. We also thank organ procurement organizations for their efforts in procuring human pancreas, as well as the organ donors and their families for their generous gifts in support of diabetes research. We thank C. Reeks for technical assistance with SIK2 Tg mice, the Histology Core Facility of Sunnybrook Research Institute for tissue processing, and K. Prentice for helpful comments. This study was funded by the Canadian Institutes of Health Research (CIHR, PJT #148931). RAS was supported by the Canada Research Chairs program (2006-2015), and the Canadian Foundation for Innovation (JELF #34677). JLR was supported by a fellowship from the Banting and Best Diabetes Centre and the CIHR (MFE #146643). JS was supported by the Japan Society for the Promotion of Science Fellows (10J00366). EM was supported by Tamarack Graduate Award in Diabetes Research from the Banting and Best Diabetes Centre.

## Author contributions

CI, JLR, EM, and QH researched data and edited the manuscript. JS and LW researched data. TK provided human islets. RAS designed the project and wrote the manuscript.

## Supplemental Data

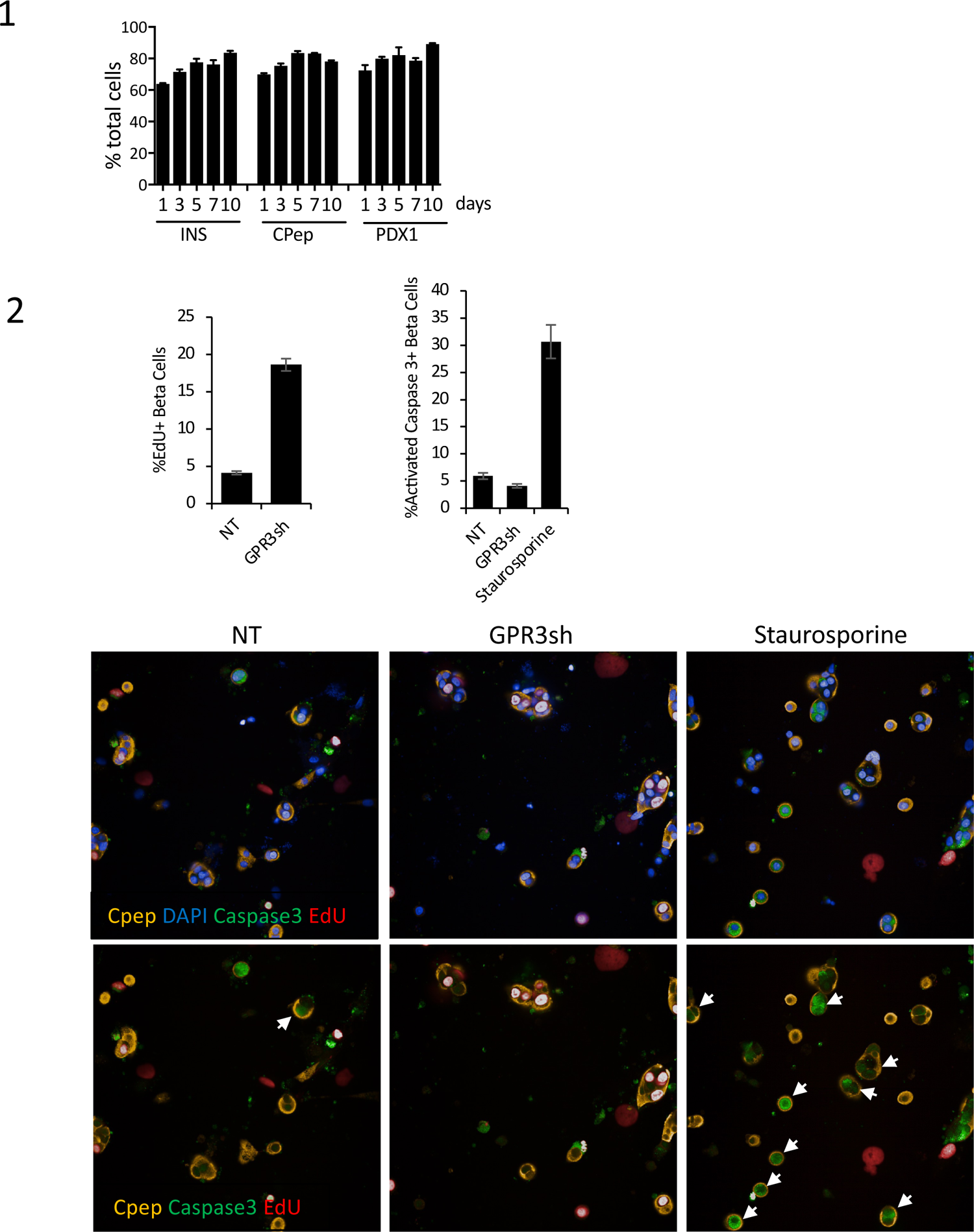
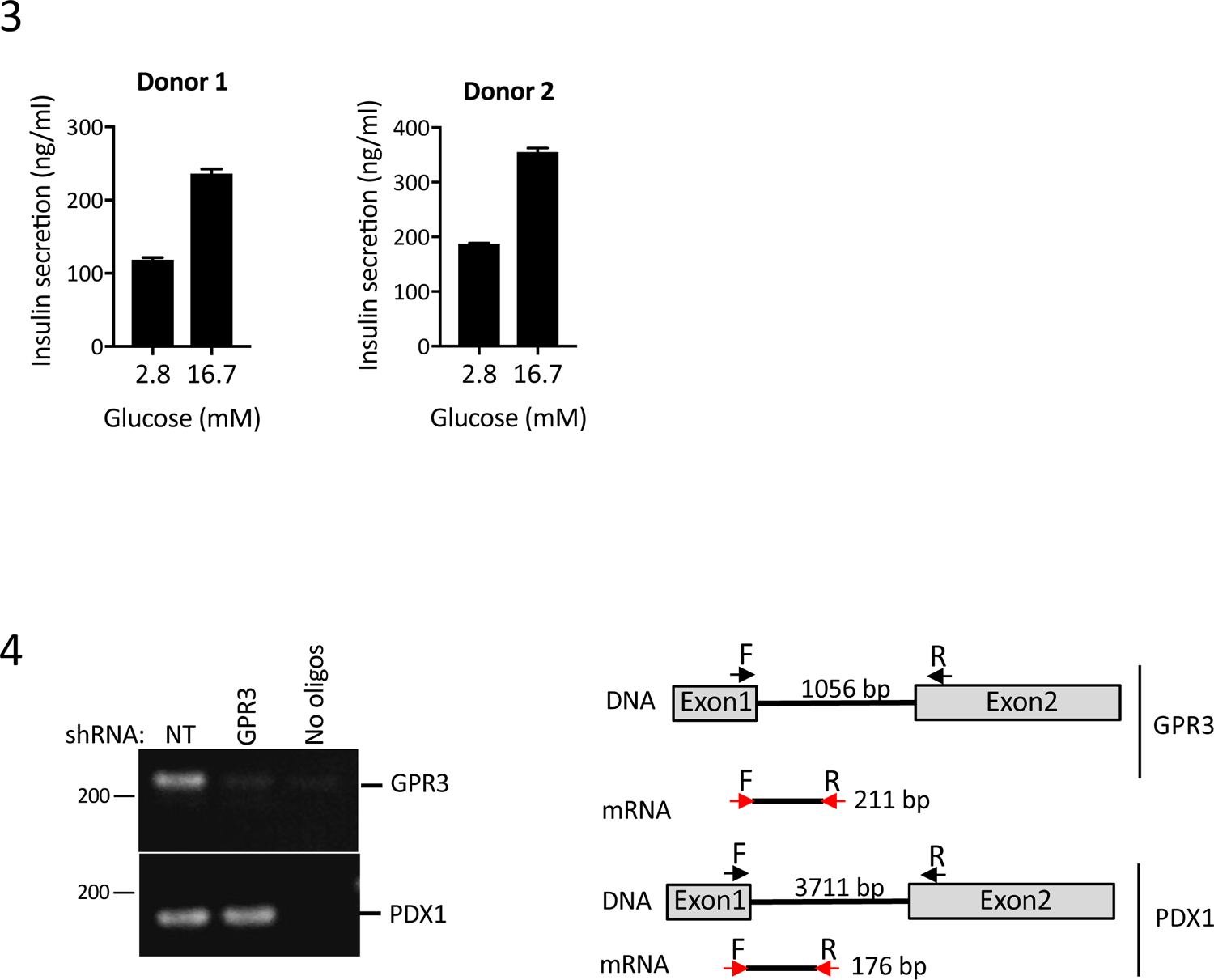
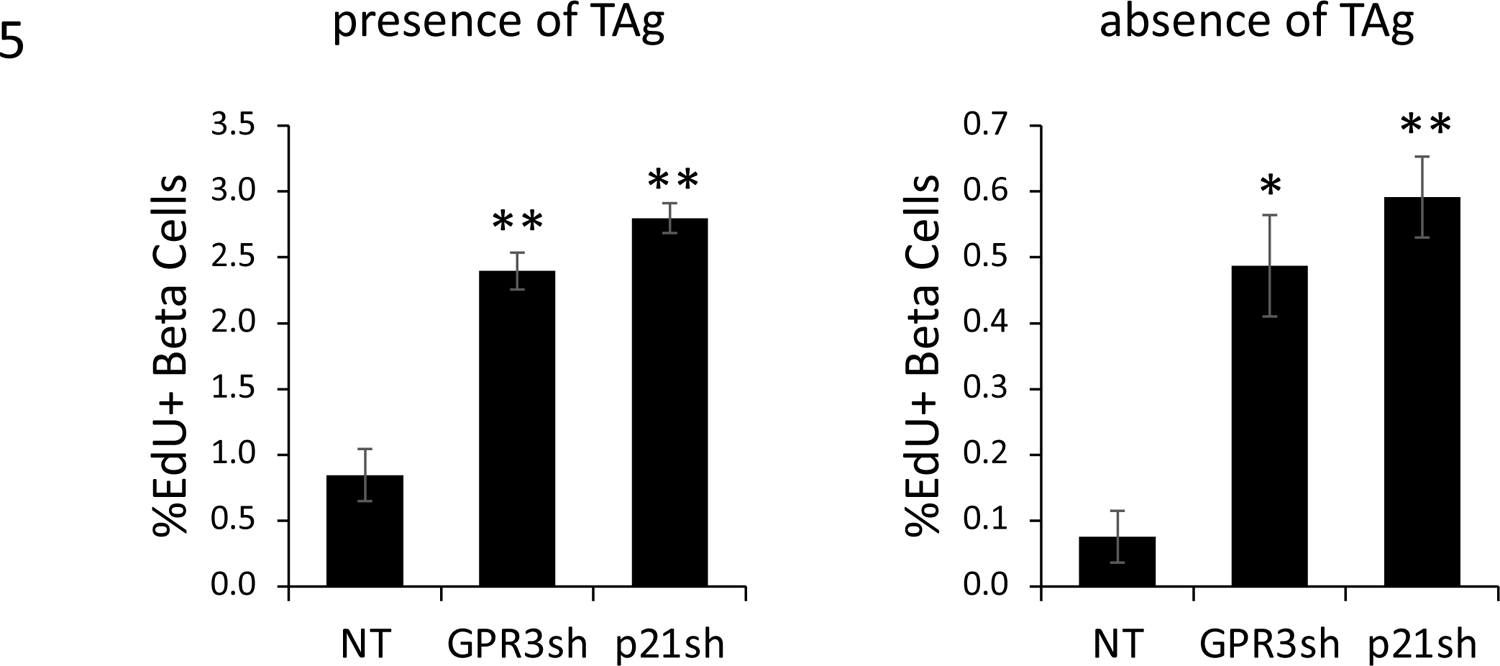
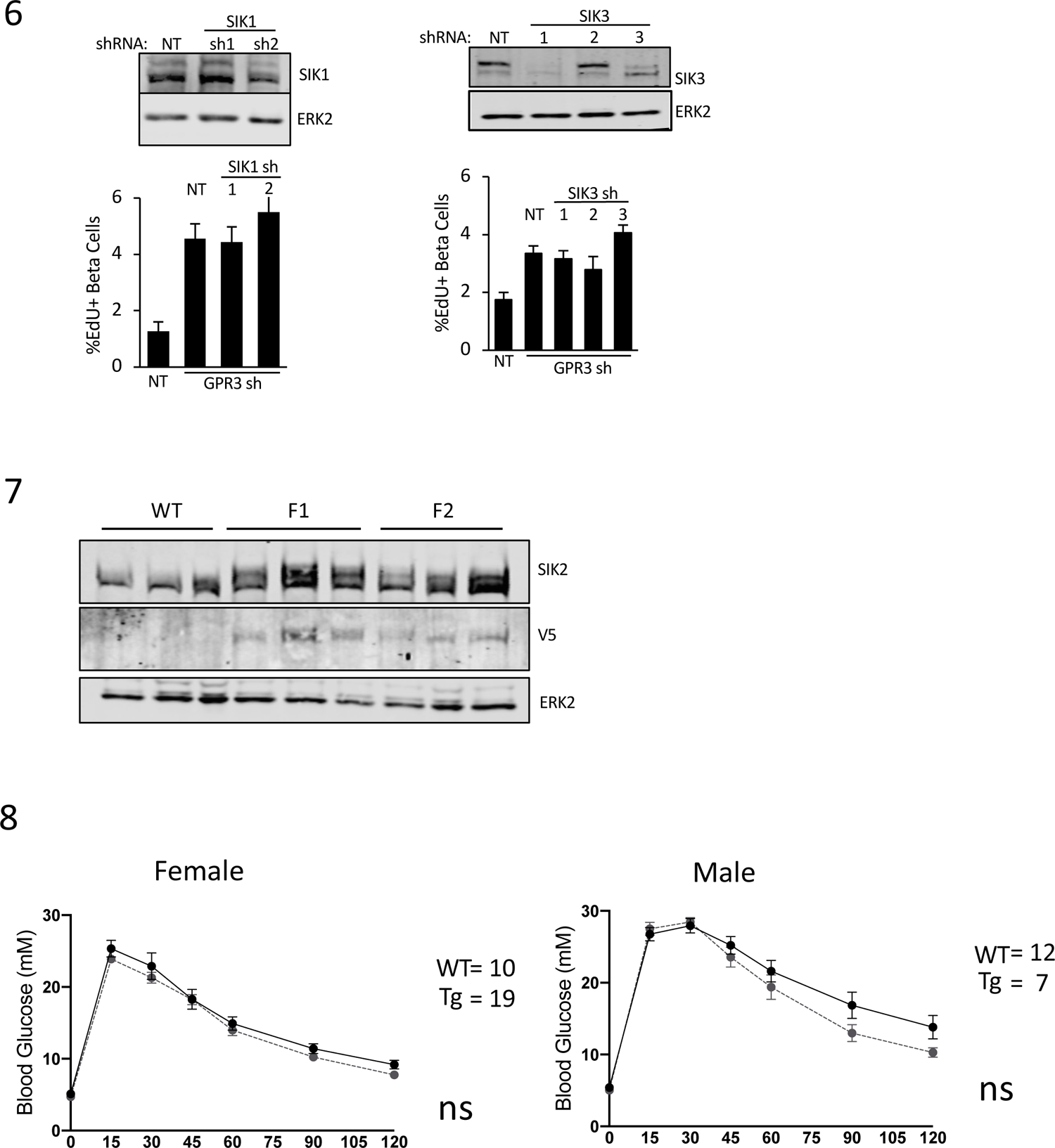
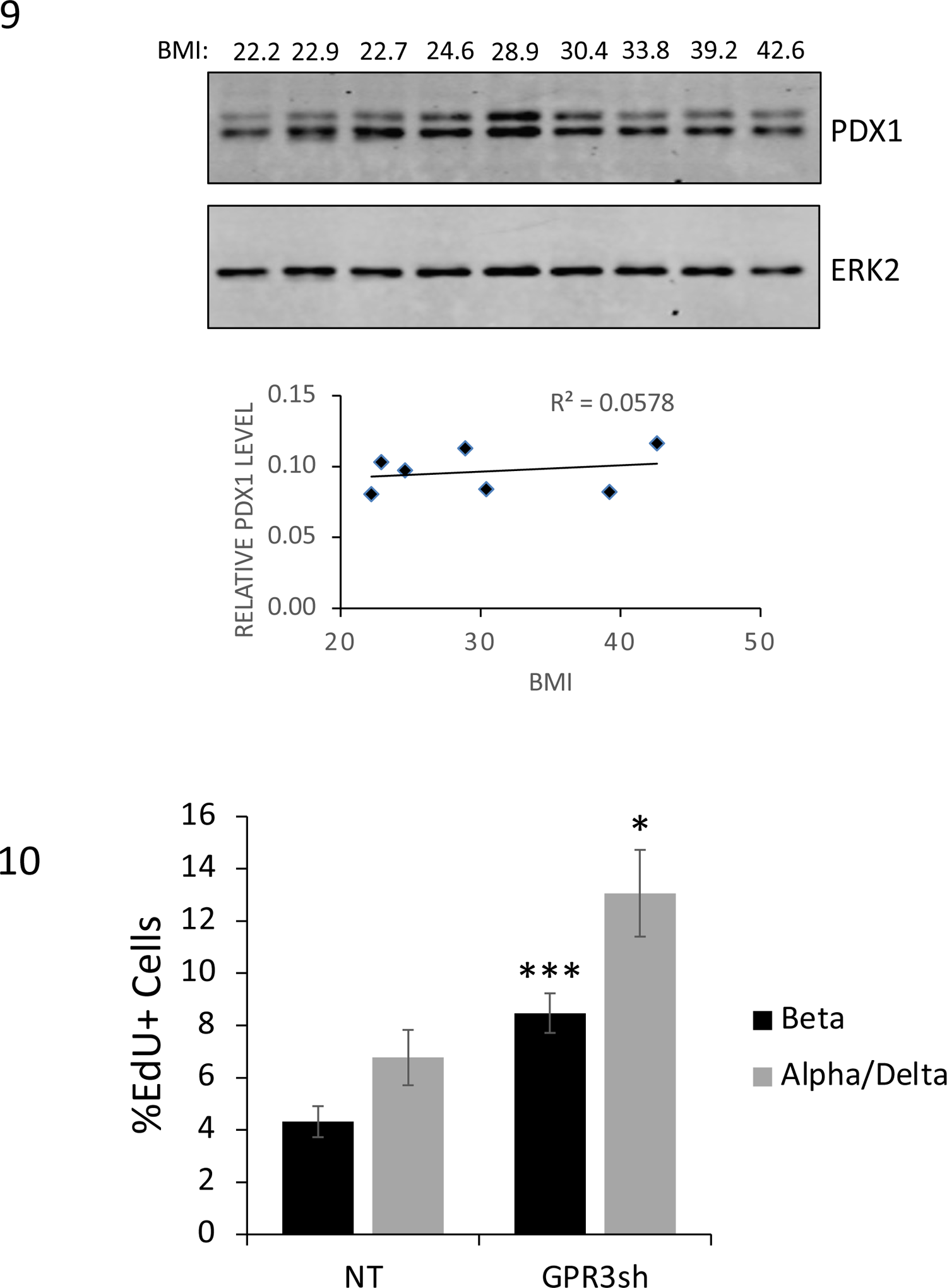
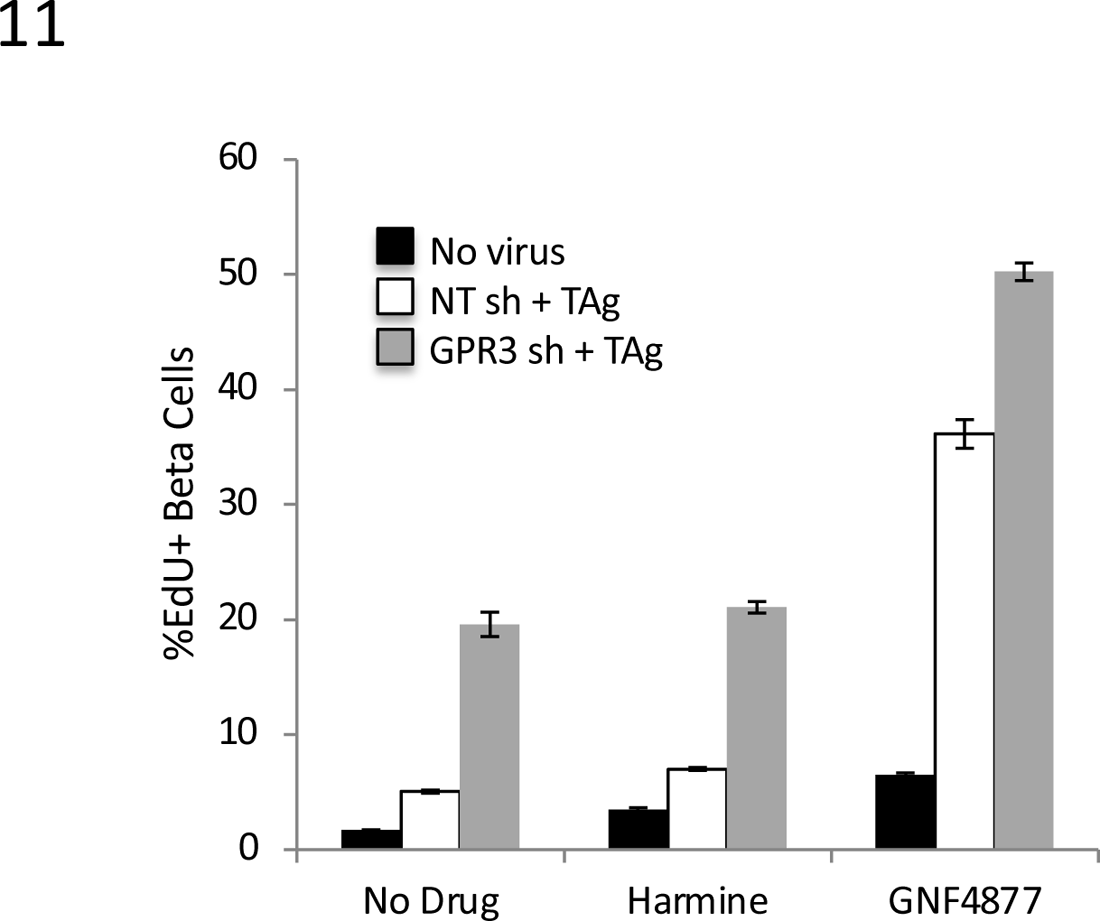
1. Barplot showing percentage of beta cells over screening assay time course, indicated by insulin (INS), C-peptide (Cpep) and PDX1 staining. n=3. 2. <u>Top</u>: % EdU+/C-peptide+ cells (left) and % activated-caspase3/C-peptide+ cells (right) in human beta cells silenced for GPR3 compared to non-targeting control (NT). Effect of the apoptotic stimulus staurosporine (40 mM) is shown (16 hr treatment). <u>Bottom</u>: Immunofluorescence micrographs of human beta cells silenced for GPR3 compared to non-targeting control (NT) stained with C-peptide (yellow), EdU (red), activated caspase-3 (green), and DAPI (blue). Effect of 40 mM staurosporine is shown. <u>NOTE</u>: activated caspase-3+ cells are not EdU+. Total number of cells counted = 2000. 3. Static GSIS data for islets from two donors that were pooled and used for GPCR screen. Treatment with low (2.8 mM) and high (16.7 mM) glucose shown. 4. (left) RT-PCR data showing loss of GPR3 mRNA in cells silenced for GPR3. PDX1 internal control shown. (right) Schematic showing RT-PCR primers spanning exons. 5. Beta cell proliferation in dissociated human islet cultures following introduction of non-targeting (NT), GPR3 or p21 shRNA in the presence (left) or absence (right) of TAg. 6. Silencing of SIK1 and SIK3 does not restore quiescence in cells silenced for GPR3. Barplots showing % proliferation with indicated combination of shRNAs and Western blots showing degree of silencing of SIK1 and SIK3 with indicated shRNAs is shown. 7. Western blots showing levels of SIK2 in WT, SIK2 Tg founder 1, and SIK2 Tg founder 2. Blot for V5 tag on transgene and ERK2 loading control shown. 8. Glucose tolerance tests in female (left) and male (right) WT or SIK2 Tg 20-week-old mice. 10. PDX1 blot for human subjects of increasing BMI. Loading control ERK2 is shown. Quantification and R^2^ value shown. 11. Beta cell or alpha/delta cell proliferation in dissociated human islet cultures following introduction of non-targeting (NT) or GPR3 shRNA in the presence of TAg. 12. Barplot showing proliferation in human beta cells following 5d treatment with pan-kinase inhibitors harmine (10 uM) or GNF4877 (2 um) alone (black bars), in combination with SV40 TAg (white bars), or in combination with SV40 TAg and GPR3 silencing (grey bars).

